## Supplementary material for "Deletion of sulfate transporter SUL1 extends yeast replicative lifespan via reduced PKA signaling instead of decreased sulfate uptake": SUL1-eLife -supplementary data

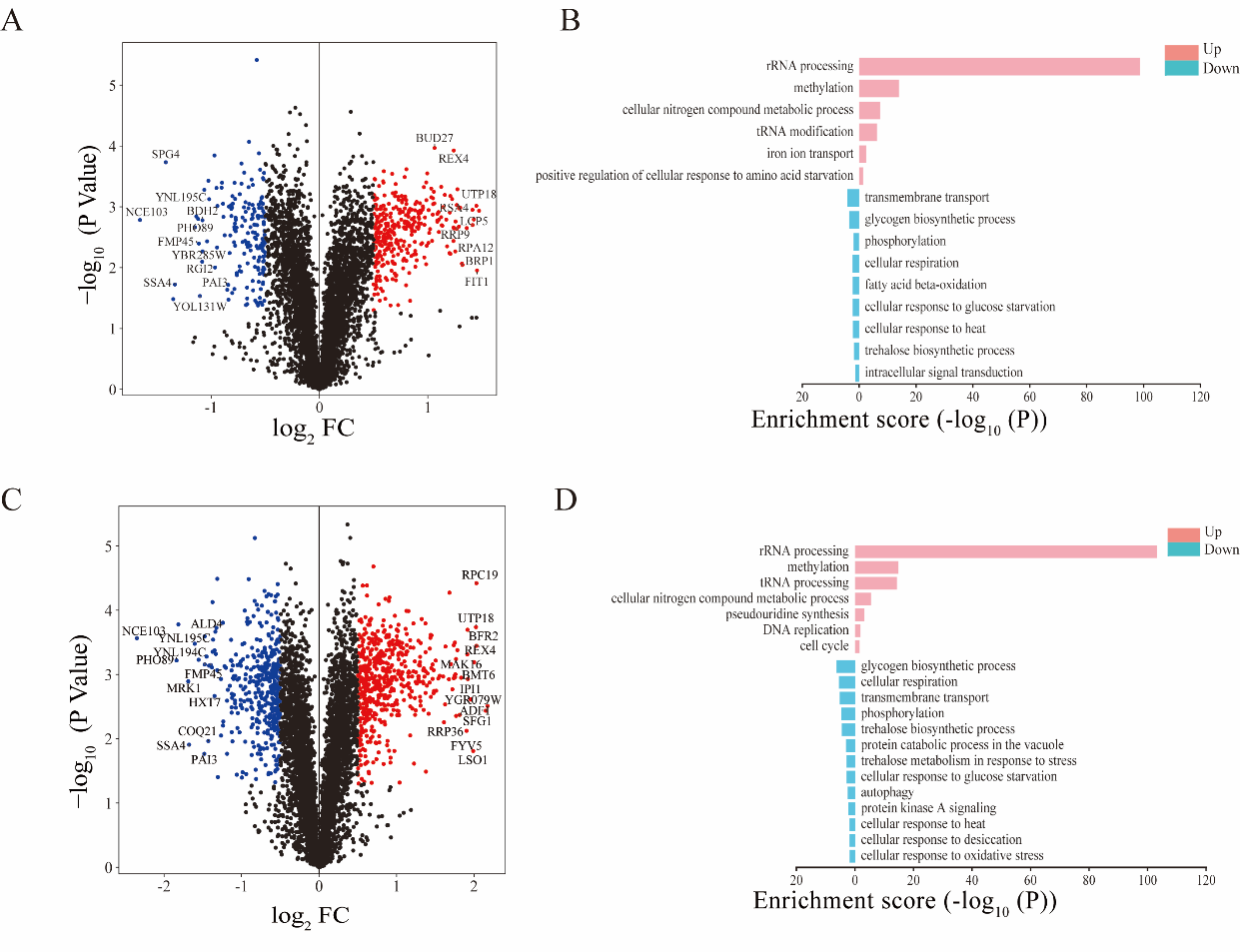

**Supplementary Figure 1.** **Transcriptome sequencing analysis were assessed in** **SUL2Δ** **VS. WT and SUL^E427Q^ VS. WT.** (A) A volcano plot illustrating the DEGs between the SUL2Δ and WT strains. Log_10_ of the *P* values plotted against the Log_2_ FC of the FPKM. (B) Enrichment analysis of biological processes associated with the DEGs identified between the SUL2Δ and WT strains. Up-regulated genes (P < 0.1, Log2 FC > 0.5) and down-regulated genes (P < 0.1, Log2 FC < -0.5) were included in this analysis. (C) A volcano plot illustrating the DEGs between the SUL^E427Q^ and WT strains. Log10 of the P values plotted against the Log2 FC of the FPKM. (D) Enrichment analysis of biological processes associated with the DEGs identified between the SUL^E427Q^ and WT strains. Up-regulated genes (P < 0.1, Log2 FC > 0.5) and down-regulated genes (P < 0.1, Log2 FC < -0.5) were included in this analysis.

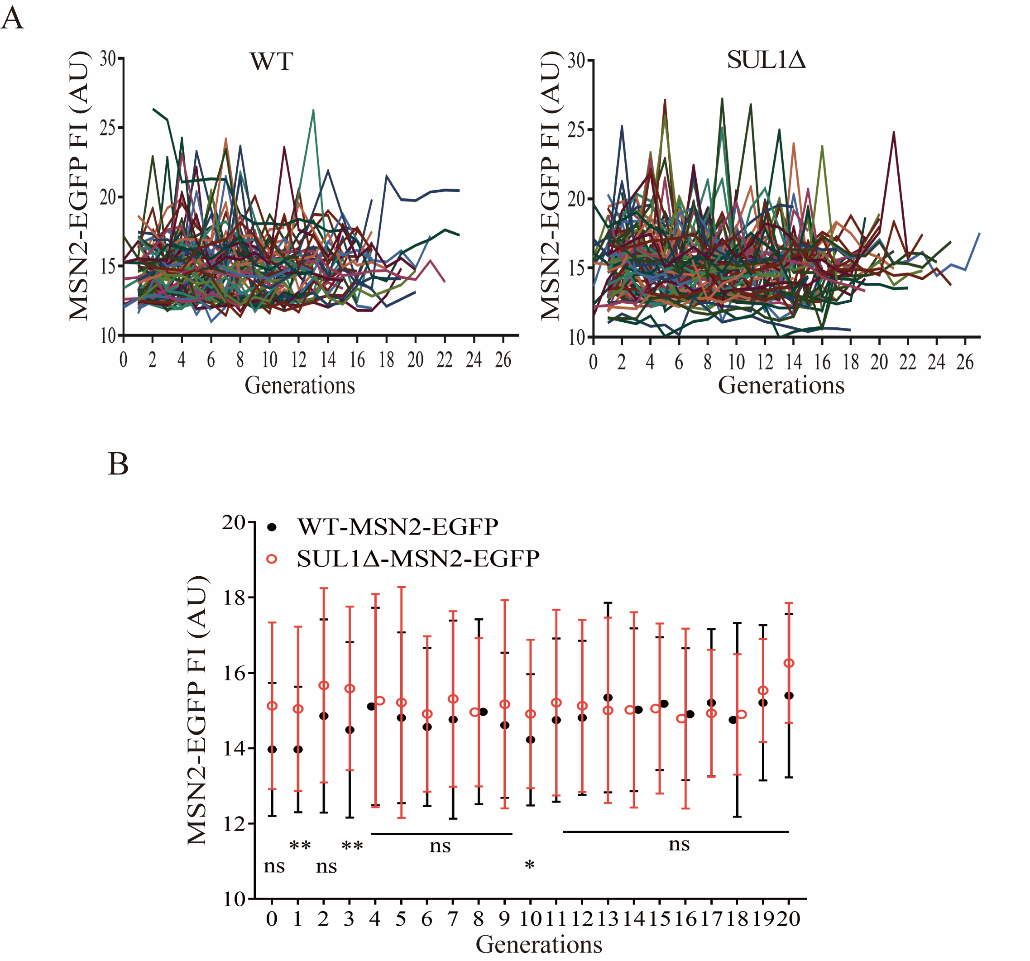

**Supplementary Figure 2.** **The expression of MSN2 protein remains stable during aging in the SUL1Δ strain.** (A) The mean fluorescence intensity (FI) of MSN2-EGFP in each generation of WT and SUL1Δ strains during RLS. (WT: n=80; SUL1Δ: n=80). (B) Compare the whole cell mean FI of MSN2-EGFP in each generation of WT and SUL1Δ strains during RLS. Bars represent mean ± SD, n=80. ns: not significant; *: *P* < 0.05; **: *P* < 0.01.

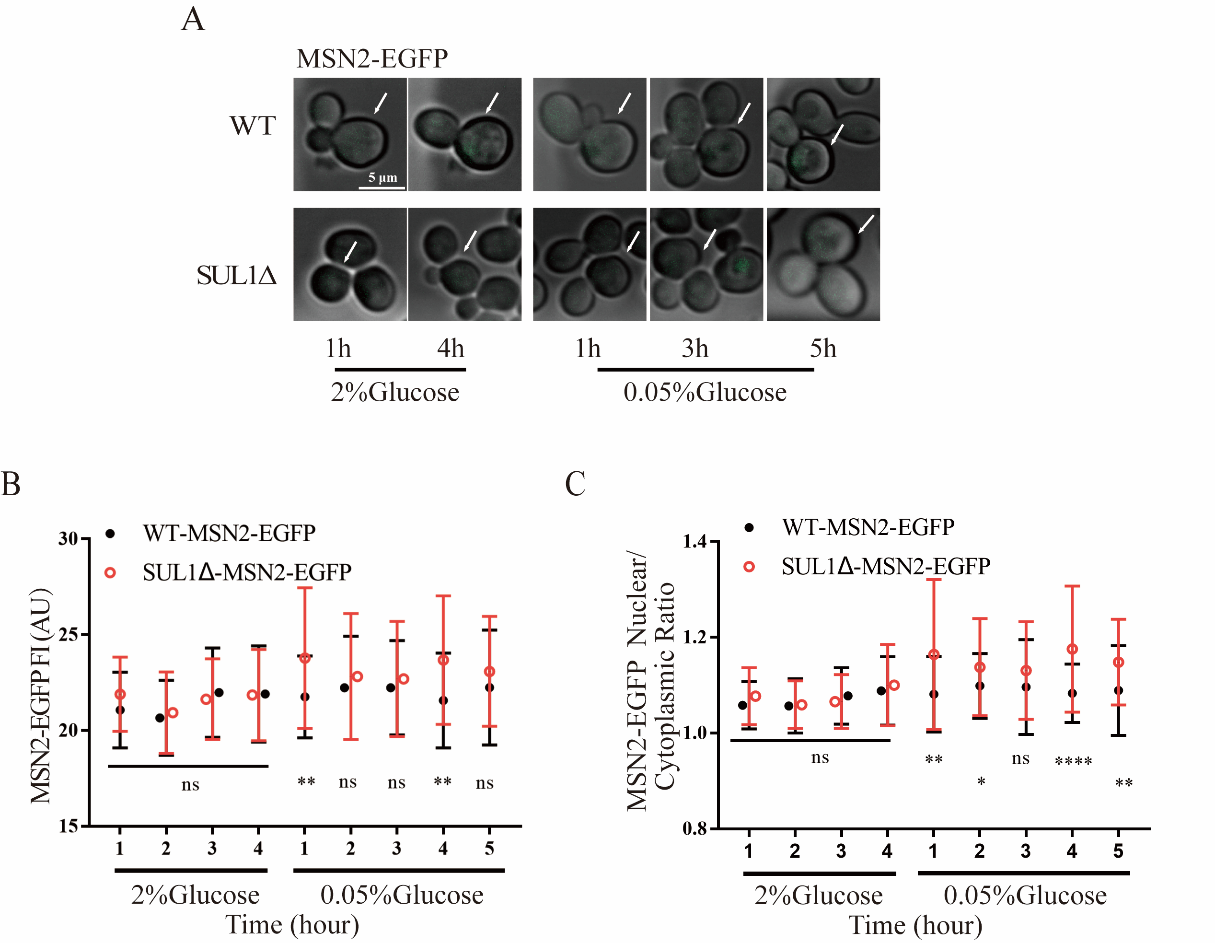

**Supplementary Figure 3.** **The nuclear translocation of MSN2 was significantly augmented upon GR stimulation.** (A) Representative time-lapse images of MSN2-EGFP in WT and SUL1Δ strains grown in complete synthetic medium (2% glucose) or in glucose restriction medium (0.05% glucose) at the indicated times. White arrows represent tracking cells. Scale bars: 5 μm. Compare the mean fluorescence intensity (B) and the nuclear/cytoplasmic mean fluorescence intensity ratio (C) of MSN2-EGFP in WT and SUL1Δ strains grown in complete synthetic medium (2% glucose) or in glucose restriction medium (0.05% glucose) at the indicated times. Bars represent mean ± SD; WT: n=43; SUL1Δ: n=40. ns: not significant; *: *P* < 0.05; **: *P* < 0.01; ****: *P* < 0.0001.

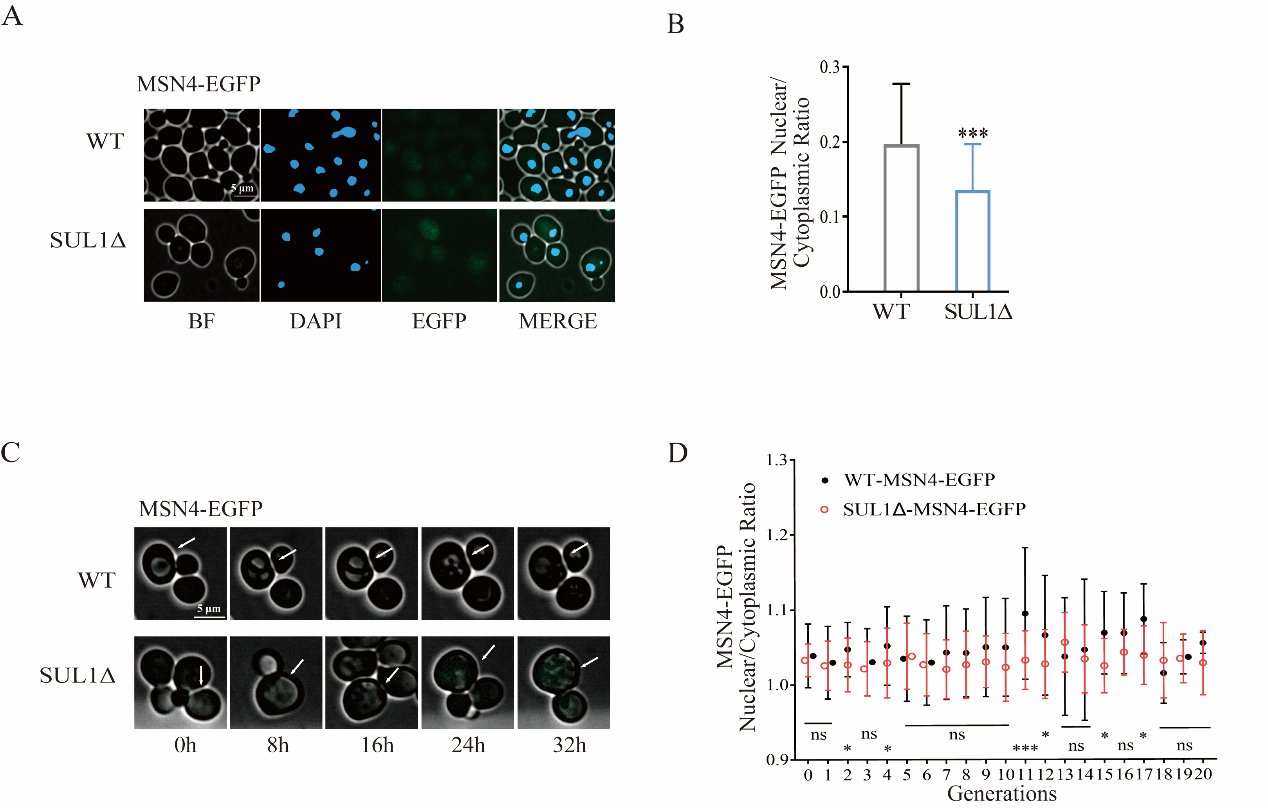

**Supplementary Figure 4. SUL1 deletion did not promote MSN4 translocation to the nucleus.** (A) Representative images of EGFP-labeled endogenous MSN4 in WT and SUL1Δ strains during the exponential growth phase. BF: bright field. Scale bars: 10 μm. (B) The ratio of the mean fluorescence intensity of MSN4-EGFP between the nucleus and the total cell. Bars represent mean ± SD, n=100. ***: *P* < 0.001. (C) Representative time-lapse images of MSN4-EGFP in WT and SUL1Δ strains. White arrows represent tracking cells. Scale bars: 5 μm. (D) Compare the nuclear/cytoplasmic mean fluorescence intensity ratio of MSN4-EGFP in each generation of WT and SUL1Δ strains during RLS. Bars represent mean ± SD, n=80. ns: not significant; *: *P* < 0.05; ***: *P* < 0.001.

Table S1. Strains Used in this Study, Related to Experimental Procedures

| **Strain** | **Genotype** |
| --- | --- |
| BY4741 sul1Δ | MATa *his3Δ1 leu2Δ0 met15Δ0 ura3Δ0 sul1Δ::KanMX* |
| BY4741 sul2Δ | MATa *his3Δ1 leu2Δ0 met15Δ0 ura3Δ0 sul2Δ::KanMX* |
| BY4741 met3Δ | MATa *his3Δ1 leu2Δ0 met15Δ0 ura3Δ0 met3Δ::KanMX* |
| BY4741 msn2Δ | MATa *his3Δ1 leu2Δ0 met15Δ0 ura3Δ0 msn2Δ::his* |
| BY4741 atg8Δ | MATa *his3Δ1 leu2Δ0 met15Δ0 ura3Δ0 atg8Δ::his* |
| BY4741 sul1Δmsn2Δ | MATa *his3Δ1 leu2Δ0 met15Δ0 ura3Δ0 sul1Δ::KanMX msn2Δ::his* |
| BY4741 sul1Δatg8Δ | MATa *his3Δ1 leu2Δ0 met15Δ0 ura3Δ0 sul1Δ::KanMX atg8Δ::his* |

Table S2. Primers Used in this Study, Related to Experimental Procedures

| **Primer** | **Sequence** |
| --- | --- |
| ko-SUL1-F | GAATATGTCACGTAAGAGCTCGACTGAATATGTGCATAATCAGGAGGATGgacatggaggcccagaat |
| ko-SUL1-R | CGGGTATATCGATATGAAAAAACGGTAAATTTGTCCCAGTTGCAGCACATcagtatagcgaccagcat |
| ko-SUL2-F | GCAGGGAATTATCTAAGATATGTCCAGGGAAGGTTATCCAAACTTTGAAGgacatggaggcccagaat |
| ko-SUL2-R | GGGATATCGATATGGAAGAAAGGTAGATTTGTTCCTGAAGCGGTATAAACcagtatagcgaccagcat |
| ko-MET3-F | CATGCCTGCTCCTCACGGTGGTATTCTACAAGACTTGATTGCTAGAGATGgacatggaggcccagaat |
| ko-MET3-R | GGTCGATCATGAATTTTGCCCTACTTTTGAGATGGGAGCATTTTATGACGcagtatagcgaccagcat |
| ko-ATG8-F | ATGAAGTCTACATTTAAGTCTGAATATCCATTTGAAAAAAGgacatggaggcccagaat |
| ko-ATG8-R | CTACCTGCCAAATGTATTTTCTCCTGAGTAAGTGACATACAcagtatagcgaccagcat |
| ko-MSN2-F | CAACTTTTATTGCTCATAGAAGAACTAGATCTAAAATGACGGTCGACCATgacatggaggcccagaat |
| ko-MSN2-R | GGGGTCTATTAAATGTCTCCATGTTTTTTATGAGTCTTGATGTGTTGCGAcagtatagcgaccagcat |
| ko-MSN4-F | TAATGCTAGTCTTCGGACCTAATAGTAGTTTCGTTCGTCACGCAAACAAGgacatggaggcccagaat |
| ko-MSN4-R | ACTTGTCATACCGTAGCTTGTCTTGCTTTTATTTGCTTTTGACCTTATTTcagtatagcgaccagcat |
| MSN2-EGFR-F | GCGATAATTTGTCGCAACACATCAAGACTCATAAAAAACATGGAGACATTggtggttctggtggtggttct |
| MSN2-EGFR-R | CAATAAGCCGTAAGCTTCATAAGTCATTGAACAGAATTATCTTATGAAGcagtatagcgaccagcattc |
| MSN4-EGFR-F | GTGACAATTTATCACAACATCTAAAAACTCACAAAAAGCACGGTGATTTTggtggttctggtggtggttct |
| MSN4-EGFR-R | CTGAGGAAGAAAGAATATTATTTCTCCGAAAACTTGTCATACCGTAGCTTcagtatagcgaccagcattc |
| ATG8-EGFR-F | AGGACGGGTTTTTGTATGTCACTTACTCAGGAGAAAATACATTTGGCAGGggtggttctggtggtggttct |
| ATG8-EGFR-R | CATTCTTATACTGGAACAATAGATGGCTAATGAGTCCCTATAATTTCGAcagtatagcgaccagcattc |
| E427Q-mutantF1 | GACGGCCAGTGAATTCATGTCACGTAAGAGCTCGACTGAATATGTGCATAAT |
| E427Q-mutantR1 | ACTACACTACTTGTCATCGTCGTCCTTGTAATCAACGTCCCATTTAGAAAAATCGGGTATATCGATA |
| E427Q-mutantF2 | TGACAAGTAGTGTAGTAGATAGTAAGTACTTTAATTACCCCCCCTGT |
| E427Q-mutantR2 | TATTCTGGGCCTCCATGTCCTCGAGCACATTAGGAGAGACAAGCCGTCCAAACT |
| E427Q-mutantF3 | GTCTCTCCTAATGTGCTCGAGGACATGGAGGCCCAGAATACCCTCCTTGACAG |
| E427Q-mutantR3 | GATTACGCCAAGCTTCAGTATAGCGACCAGCATTCACATACGATTGA |
| E427Q-mutantF427 | GTTGTCCCTGACCAACAACTTATTGCGATTG |
| E427Q-mutantR427 | CAATCGCAATAAGTTGTTGGTCAGGGACAAC |
| q-SUL1-F-1 | TGAGGTTTTACCAGCCCCAG |
| q-SUL1-R-1 | TGCCTTGAGCAGTTCACGTA |
| q-SUL2-F-1 | CCGACCAAGAATTGATTGCT |
| q-SUL2-R-1 | AAGAATGCGCCGGTCAAACA |
| q-MET3-F | GGTCCATACGATGCTCAAGA |
| q-MET3-R | CTTGTTTTGGTCTTGGTGGG |
| q-MSN2-F | CCAAGTCAGCAATTACAGCA |
| q-MSN2-R | CGGCAGCATATCGTTCTTGG |
| q-MSN4-F | TCGCGACGCAAGAAGATACA |
| q-MSN4-R | GTTGATCCACGCCTCTCAGT |
| q-ATG8-F | AGTTCCTGCTGACCTTACCG |
| q-ATG8-R | ACCCGTCCTTATCCTTGTGTT |
| q-HSP12-F | ACATCACTGACAAGGCCGAC |
| q-HSP12-R | GCGGCTCCCATGTAATCTCT |
| q-ALD3-F | ACATCTTCTGAGCAACGTGGTA |
| q-ALD3-R | CCGCCCCCGCATAGTATCTT |
| q-RTN2-F | GTTCACGAGTTTTGCGGTGG |
| q-RTN2-R | TTCTGTCCTCTAACGCAGGC |
| q-SIP18-F | GGAAAGAACGCCAAATCCTCC |
| q-SIP18-R | TCCAATCGTTCGCAATTCCTC |
| q-CTT1-F | GCAATTCCACGTCTTGTCGG |
| q-CTT1-R | AGTTGCTTGTTCGGGTGTCA |
| q-TPS1-F | GCCGTACCCATCTTCCTGAG |
| q-TPS1-R | GGTTTGCCTCGTTGTATGCC |
| q-PNS1-F | TATGCACGGTAGGCGGATTC |
| q-PNS1-R | GGATGGCAGCATTGGTGTTC |
| q-ACT1-F-1 | CGTTTCCATCCAAGCCGTTT |
| q-ACT1-R-1 | ACCGGCCAAATCGATTCTCAA |
